# High-performance machine learning for peptide classification from nanopore translocation events, leveraging event kinetics and duration filtering

**DOI:** 10.1101/2025.08.21.671556

**Authors:** Jennifer M. Colby, Bryan A. Krantz

## Abstract

Understanding single-molecule translocation dynamics through biological nanopores is fundamental to advancing next-generation biosensing and sequencing technologies. Here, using the anthrax toxin protective antigen nanopore, we describe a high-performance machine learning (ML) framework for classifying a diverse series of guest-host peptides based on individual translocation events. The approach leverages carefully engineered, event-level biophysical features extracted from either scaled current and conductance state sequences. Through systematic UMAP analysis of this feature space, we reveal that filtering away the shortest events effectively enriches the dataset with more discriminative longer events, leading to improved classification. Various deep learning (DL) and traditional ML architectures, including convolutional neural networks (CNN), temporal convolutional networks (TCN), and eXtreme Gradient Boosting (XGBoost), were investigated. The dual-input CNN-Dense model, which utilized current sequences and features, achieved strong classification performance (accuracy ∼0.80). However, the most robust classification was achieved with XGBoost acting solely on the engineered feature set, demonstrating superior performance (accuracy ∼0.90). This ML approach provided a significant computational advantage in both training and inference over DL models. Notably, these models consistently discriminated between peptides differing only in backbone stereochemistry, highlighting the exquisite sensitivity of the nanopore to subtle conformational dynamics. These findings underscore that carefully engineered event-level features, particularly from longer translocations, combined with efficient tree-based models, offer a highly effective and computationally favorable strategy for high-fidelity peptide classification for biosensing applications.

## Introduction

The rapid and accurate detection and characterization of biomolecules, particularly peptides and proteins, represent a critical and often unmet challenge across diagnostics, drug discovery, and fundamental biological research. Peptide biomarkers, indicative of various disease states including heart disease, infectious diseases, and cancer (1–3), offer immense potential for timely and precise diagnosis. However, analyzing complex peptide mixtures *in situ*, often at low concentrations, demands analytical technologies with unparalleled sensitivity and specificity—capabilities that remain challenging for many conventional methods. Single-molecule nanopore biosensing provides a transformative approach (4), enabling the detection and characterization of individual molecules as they traverse a nanometer-scale pore by measuring picoamp-scale modulations in ionic current. Beyond simple presence/absence detection, nanopore technology holds the promise to revolutionize biopolymer analysis, including the ambitious goal of direct, high-throughput peptide and protein sequencing, a major frontier where robust, widely applicable methods are still lacking (5–7).

Nanopore biosensors consist of a membrane-embedded pore separating two electrolyte-filled compartments. Under an applied driving force (either a voltage or proton gradient), biomolecules are directed through the pore, generating unique current signatures dependent on their size, shape, and chemical properties. Biological protein nanopores, such as those formed by transmembrane proteins inserted into lipid bilayers, offer exquisite control over pore geometry and molecular interactions (8–14). Single-channel recordings capture the dynamic, stochastic interactions of individual molecules with the nanopore in high-resolution ionic current traces. Extracting the maximum analytical information from these complex, high-dimensional translocation event streams is the central bottleneck preventing the full realization of nanopore technology’s potential for analyzing complex biological samples.

Among the most promising biological nanopores for polypeptide analysis is the anthrax toxin protective antigen (PA) channel from *Bacillus anthracis* (15) (Fig. 1A). As a natural protein translocase, PA possesses unique structural/functional features, which are highly advantageous for peptide biosensing and sequencing. It is remarkably robust, facilitates highly processive polypeptide translocation driven by an applied voltage (16) or proton gradient (17) with distinct kinetic substeps, and utilizes multiple internal ‘peptide-clamp’ sites (9, 12–14, 18, 19) to engage heterologous polypeptides (20) without the need for cumbersome tags like DNA handles (18, 19, 21, 22). Importantly, polypeptide translocation through the PA pore generates rich, multi-state current signatures (18), often involving multiple discrete partially blocked sub-conductance intermediates in addition to the fully blocked and open states. These distinct, peptide-dependent kinetic and conductance characteristics are information-rich readouts that can serve as powerful discriminatory features for peptide identification and classification in complex mixtures.

**Fig. 1.**
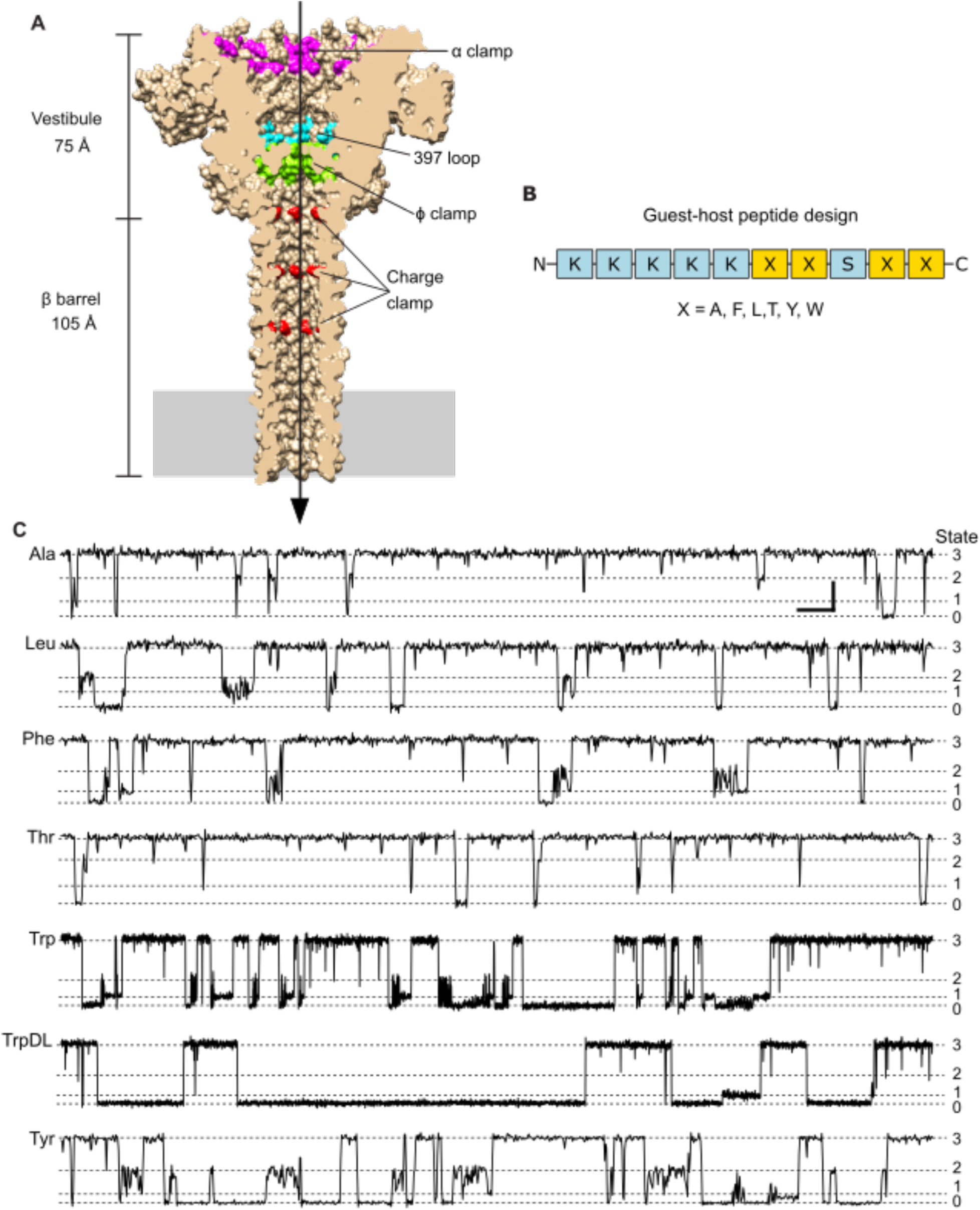
PA nanopore peptide biosensor. **(A)** Sagittal section of the anthrax toxin PA_7_ nanopore cryo-EM structure (PDB: 3J9C) (8) rendered in Chimera (37) as a molecular surface. Peptide clamps and loop active sites are colored and labeled: α clamp (magenta), 397-loop (cyan), ϕ clamp (green), and charge clamp (red). Overall scale of the upper vestibule and elongated lower β barrel are indicated. Narrowest point, at the ϕ clamp, has a luminal diameter of 6 Å. Direction of translocation from *cis* to *trans* is indicated by an arrow. Membrane bilayer position is indicated with a solid gray rectangle. **(B)** Guest-host peptide design schematic for the 10-residue guest-host peptide with general sequence, KKKKKXXSXX, with guest residue (X). **(C)** Representative current versus time records of guest-host peptide translocations carried out at 70 mV (*cis* positive) in symmetric succinate buffer, 100 mM KCl, pH 5.6. To the left are guest-host peptide names. In general, this nanopore-peptide system populates multiple discrete conductance state intermediates (indicated by dashed lines), which are enumerated by state on the far right: fully blocked (state 0), partially blocked intermediates (state 1 and state 2), and fully open (state 3). Scalebar at upper right for guest-host Ala, Leu, Phe, Thr, and Tyr peptides represents 2 pA by 100 ms. For guest-host Trp and TrpDL peptides, the scalebar represents 2 pA by 500 ms due to their characteristically longer events.

Furthermore, the fine structure and temporal dynamics of these multi-state transitions encode information potentially sufficient to resolve amino acid sequence, presenting a unique opportunity for developing label-free, peptide sequencing capabilities.

Analyzing the massive, complex, and often noisy datasets generated by multi-state nanopore systems requires advanced computational approaches. Traditional manual or simple threshold-based analysis methods are fundamentally inadequate for extracting the full information content from nuanced multi-state kinetics or dissecting complex mixtures of analytes. Machine learning (ML) and deep learning (DL) offer powerful, modular, and adaptable tools uniquely suited to identify subtle patterns in translocation state sequences and correlate them with computed biophysical features (23–27). While ML/DL has been applied to nanopore data, its application to analyzing real-world, multi-state peptide translocation data, either for precise peptide classification in mixed samples or for determination of sequence information, remains largely an underdeveloped area.

Here, leveraging anthrax toxin’s PA nanopore, we describe a high-performance ML framework for classifying a diverse series of guest-host peptides based on individual translocation events.

## Results

### Diverse guest-host peptide translocation events via PA nanopores

Previous investigations extensively characterized the broad ensemble properties and single-channel dynamics of the guest-host peptide series translocating through the anthrax toxin protective antigen (PA) nanopore (18, 21). Building upon this foundation, we sought to explore whether the intrinsic dynamic/kinetic properties of these peptides, as observed during single-channel translocation events, could serve as information-rich signatures for classification. Specifically, we aimed to assess if the PA nanopore, when coupled with powerful computational ML/DL methods, could reliably distinguish peptides based on subtle sequence differences that are challenging to discern by conventional analysis. This study thus assesses the PA nanopore’s potential as a sophisticated peptide biosensor.

Relative to other protein nanopores commonly utilized in biosensing and nucleic acid sequencing, the PA nanopore **(Fig. 1A)** possesses a significantly longer lumen. This architecture, however, features numerous active site clamps (e.g., α-clamp, ϕ-clamp, and charge clamp) and loop regions (e.g., 397-loop), specifically evolved to processively translocate large (∼100 kDa) proteins, and, at nanomolar concentrations, shorter peptides. The 10-residue guest-host peptide series, with a general sequence of KKKKKXXSXX, was initially designed to systematically probe differences in binding, translocation, and dynamics based on the guest residue (X) **(Fig. 1B)**. Our guest-host panel included peptides with standard natural ʟ-stereochemistry for guest residues Ala, Leu, Phe, Thr, Trp, and Tyr. To investigate the sensitivity of the nanopore to stereochemical variations, a seventh peptide, called guest-host TrpDL, was included; it shared the same amino acid sequence as guest-host Trp but featured an alternating pattern of ᴅ- and ʟ-stereoisomers along its backbone.

For this assessment of the PA nanopore as a biosensor platform, we focused on collecting extensive single-channel translocation event streams via planar lipid bilayer electrophysiology. The data acquisition and processing workflow is shown in **Fig. S1A**. All analyses presented herein utilized data acquired under a consistent 70 mV driving force (*cis* positive). This potential strongly favors complete translocation events, which is critical for consistent feature extraction as it ensures the peptide fully interacts with the entire nanopore rather than more superficially engaging the entrance, as might occur at lower potentials.

Furthermore, the signal-to-noise ratio is inherently higher at larger potentials, providing additional support for this selection. A comprehensive breakdown of the total recording times per peptide in our complete dataset is provided in **Table S1**.

Across samples of translocation event streams for the seven guest-host peptide classes, four discrete conductance states are consistently observed, which are enumerated as states 0-3 **(Fig. 1C)**. However, the translocation event dynamics, as characterized by current blockade patterns and durations, cover a broad range. Qualitatively, events for guest-host Trp and guest-host TrpDL exhibit noticeably longer durations compared to the other five peptides. In contrast, peptides with smaller guest residues, such as guest-host Ala and guest-host Thr, display rapid dynamics that are often difficult to distinguish reliably by visual inspection alone. For peptides like guest-host Leu and guest-host Phe, which are similar on hydrophobicity scales, subtle visual differences in their translocation dynamics exist but are similarly challenging to resolve.

The aromatic guest residues Phe and Tyr present event lengths and flickering dynamics that are visually analogous and difficult to discriminate. Therefore, these qualitative observations, particularly the subtle or complex nature of visual distinctions between peptide classes, directly suggest that advanced ML/DL methods can be effectively exploited to robustly classify these peptides, even from individual single-channel translocation events.

### Discriminatory potential of engineered features is event length dependent

Achieving robust peptide classification from individual translocation events necessitated a comprehensive and generalized feature engineering approach. To ensure high-fidelity state assignments critical for feature extraction, raw current records were state-labeled using the ‘Single-Channel Search’ routine in CLAMPFIT, providing expert-validated assignments. For our specific four-state event streams (18), a consistent enumeration scheme was used: state 0 for fully blocked, states 1 and 2 for intermediate blockades (closest to state 0 and state 3, respectively), and state 3 for the fully open state **(Fig. 1C)**. From these labeled records, both conductance state sequences and scaled current sequences were segmented for each translocation event, from which a rich feature set (including scalar, vector, and matrix features) was calculated (see Materials and Methods where all features are delineated) **(Fig. S1B)**. While the segmentation process offers filtering and baseline correction capabilities, these were not employed for the current datasets. However, we critically explored the impact of minimum event duration as a preprocessing filter. This filter initially aimed to remove spurious, very rapid blockade dynamics, which are sometimes attributed to water structure formation (’wetting’ and ‘dewetting’) around the ϕ clamp (28). Beyond this initial rationale, it became evident through preliminary analyses on simulated data that shorter events inherently possessed less discriminatory information, analogous to attempting to classify images with significantly fewer pixels. This observation motivated a systematic investigation into the effect of minimum event duration on feature discriminability. The main practical downside to using a minimum event duration filter is that translocation event data are exponentially distributed, and higher filtering values for this parameter could remove large numbers of translocation events from the dataset **(Table S2)**.

To qualitatively and quantitatively assess the impact of different minimum event duration thresholds on the discriminative power of the extracted feature sets, Uniform Manifold Approximation and Projection (UMAP) dimensionality reduction was employed to generate 2D cluster representations. For events filtered at a minimum duration of 5 ms, UMAP analysis **(Fig. 2A)** resulted in poor clustering performance. Visually, peptide classes remained largely intermixed, with few well-structured, distinct clusters and numerous ‘stray’ points dispersed throughout the 2D projection. In contrast, increasing the minimum event-length filter to 20 ms visibly improved clustering in the UMAP analysis **(Fig. 2B)**. While some intermixing persisted and a smaller fraction of stray points remained, distinct clusters became apparent for several peptides, notably Phe, Thr, Trp, and Tyr. Interestingly, despite its strong classification performance in subsequent ML/DL models (as shown later), the events for guest-host TrpDL in the 20 ms UMAP embedding presented as several smaller, somewhat dispersed clusters, suggesting inherent sub-populations or more complex relationships not fully captured by the 2D projection. To quantify these visual observations, clustering metrics Adjusted Rand Index (ARI) and Normalized Mutual Information (NMI) were computed by applying K-Means clustering (K=7, for seven peptide classes) to the UMAP embeddings and comparing the resulting clusters to the known peptide labels. For data filtered at 5 ms, the ARI was 0.0356 and NMI was 0.0402.

**Fig. 2.**
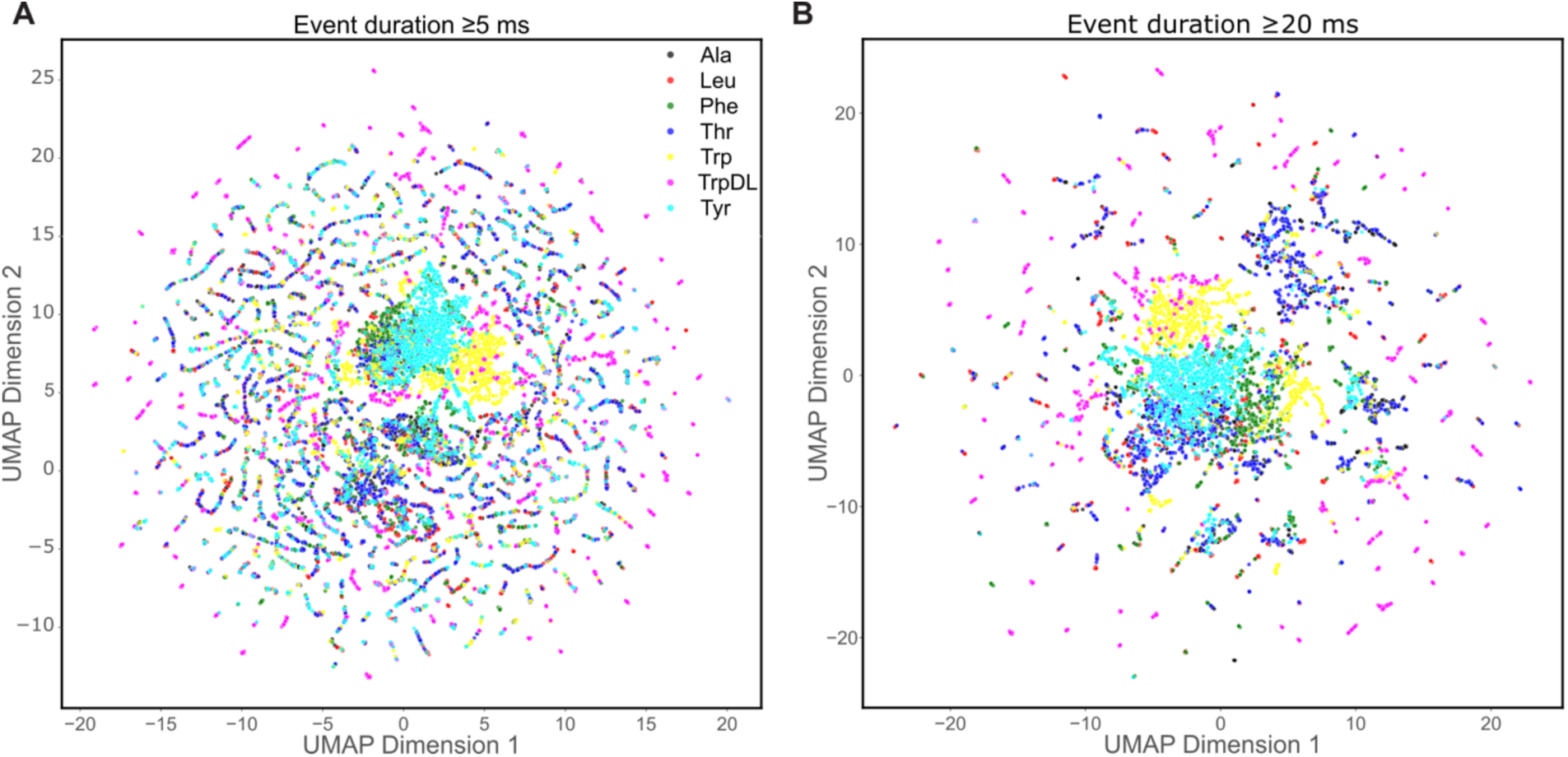
Discriminative potential of extracted features at different minimum event durations. UMAP clustering analysis of event-level features. Data points represent individual translocation events, colored by their corresponding guest-host peptide identity: Ala (black), Leu (red), Phe (green), Thr (blue), Trp (yellow), TrpDL (magenta), and Tyr (cyan). **(A)** UMAP embedding of events filtered at a minimum duration of 5 ms (n_neighbors = 15, min_dist = 0.5). Peptide classes are more intermixed with less clear separation, indicating more limited discriminative power of features for very short events (ARI: 0.0356; NMI: 0.0402). **(B)** UMAP embedding of events filtered at a minimum duration of 20 ms (n_neighbors = 15, min_dist = 0.5). In contrast, events filtered at 20 ms show visibly improved clustering for several guest-host peptides (e.g., Phe, Thr, Trp, Tyr), albeit TrpDL is more spread out and peripheral. While some scattered points remain, suggesting inherent event variability or limitations of the 2D projection, the overall separation is markedly enhanced (ARI: 0.0887; NMI: 0.1282).

However, for data filtered at 20 ms, the ARI increased to 0.0887 and the NMI to 0.1282. The substantially higher ARI and NMI values for the 20 ms filtered data strongly indicate a greater alignment between UMAP-derived clusters and true peptide identities at longer event durations. These results collectively demonstrate that the engineered feature sets gain significant discriminative power by excluding very short events, albeit with the inherent trade-off of reducing the total number of analyzed events **(Tabel S2)**.

### Performance of DL classification models

The core design approach for DL-based peptide classification from individual translocation events involved a branched, dual-input neural network architecture **(Fig. 3A, S1C)** (29). In this configuration, either the conductance state sequences (S) or the raw current sequences (C) from translocation events served as input to a multi-layered CNN or TCN branch. Simultaneously, the corresponding event-level features (F), extracted during preprocessing, were fed into a separate fully connected (Dense) network. The outputs of these two parallel branches were then concatenated before leading to a final classification layer. Based on this design pattern, three distinct configurations were assessed: TCN-Dense (S+F), CNN-Dense (S+F), and CNN-Dense (C+F). Given that the discriminative power of the engineered features was shown to be dependent on the minimum event duration parameter during preprocessing **(Fig. 2)**, these models were subjected to an initial, single-replicate performance scan across a range of minimum event duration values (5, 7.5, 10, 12.5, 15, and 20 ms) **(Table S3)**. This preliminary scan revealed a consistent trend across all DL models: increasing the minimum event duration to exclude shorter, less information-rich events significantly improved classification performance. Finalized, comprehensive performance metrics (mean ± standard deviation, N=5 independent training runs) were compiled for these models at the two extreme minimum event duration cutoffs (5 and 20 ms) **(Table 1)**.

**Fig. 3.**
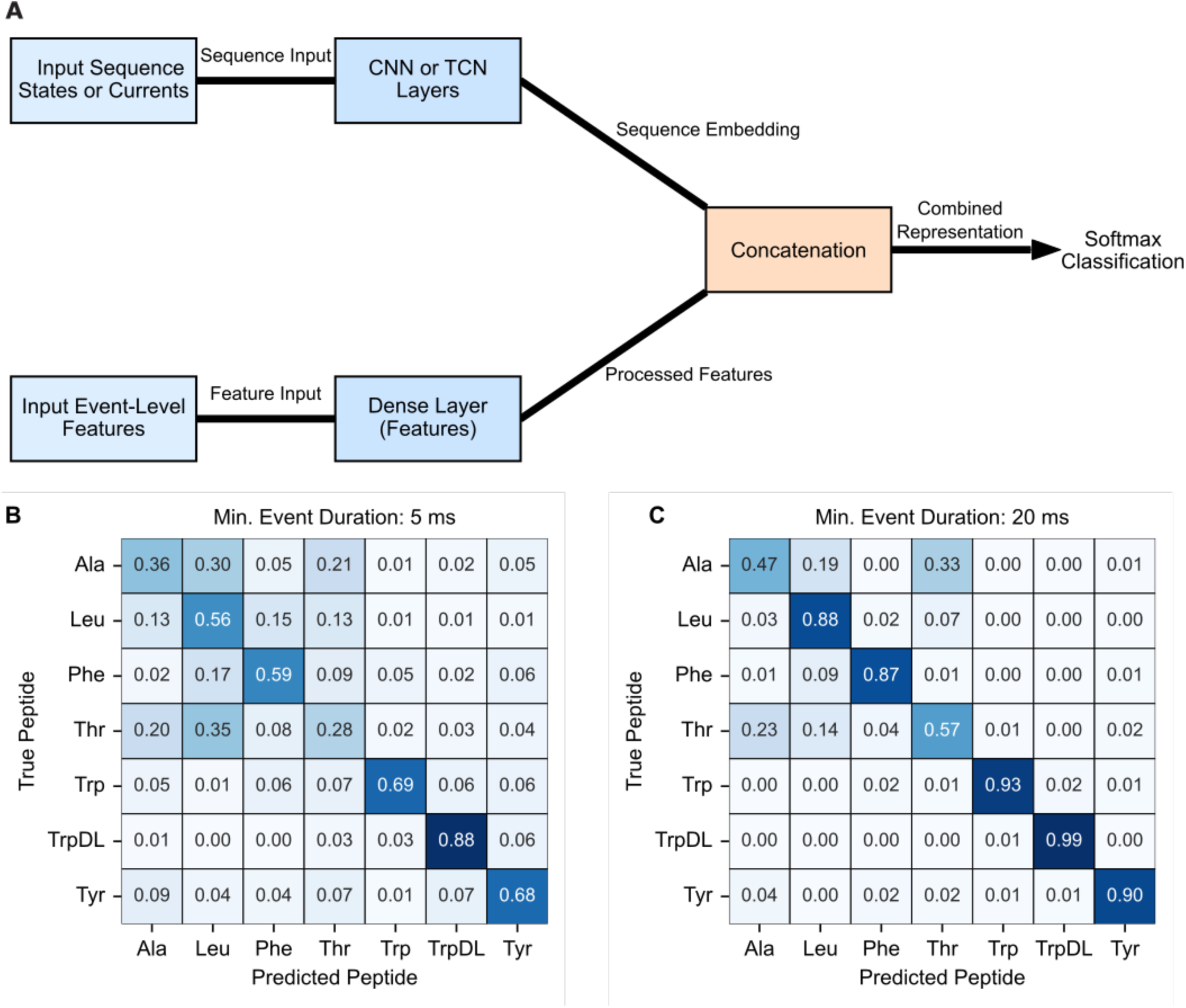
DL-based classification of guest-host peptide translocation events. **(A)** General dual-input neural network architecture as a block diagram illustrating the common design pattern for the DL classifiers. Sequence data (either conductance states (S) or raw current (C)) are processed by a multi-layered CNN or TCN branch. Concurrently, event-level features (F) are processed by a fully connected (Dense) branch. The outputs of these two branches are concatenated and fed into a final Dense layer for classification. This architecture was the basis for the TCN-Dense (S+F), CNN-Dense (S+F), and CNN-Dense (C+F) models. **(B)** Normalized confusion matrix displaying the per-class prediction accuracy of the CNN-Dense (C+F) model on the test set for a minimum event duration of 5 ms. Rows represent true peptide labels, and columns represent predicted labels. Values indicate the proportion of events from a given true class that were predicted as each class. **(C)** Normalized confusion matrix for CNN-Dense (C+F) at 20 ms minimum event duration, which is plotted as in panel B. Note the improved diagonal elements (correct predictions) and reduced off-diagonal elements (misclassifications) compared to the shorter minimum event duration predictions in panel B.

**Table 1.**
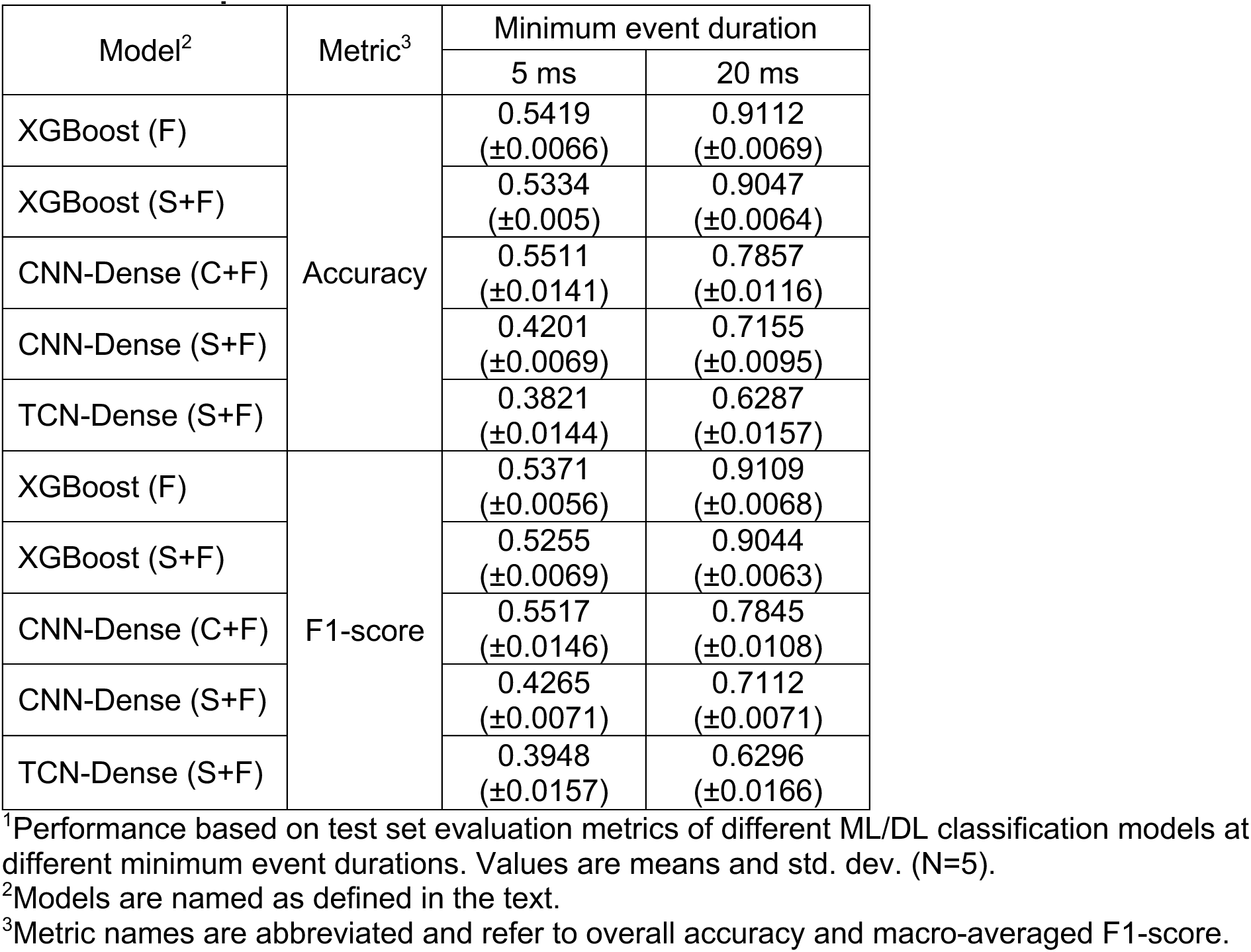
Model performance metrics at two minimum event duration extremes.^1^.

| Model<sup>2</sup> | Metric<sup>3</sup> | Minimum event duration |  |
| --- | --- | --- | --- |
|  |  | 5 ms | 20 ms |
| XGBoost (F) | Accuracy | 0.5419<br>($\pm 0.0066$) | 0.9112<br>($\pm 0.0069$) |
| XGBoost (S+F) | | 0.5334<br>($\pm 0.005$) | 0.9047<br>($\pm 0.0064$) |
| CNN-Dense (C+F) | | 0.5511<br>($\pm 0.0141$) | 0.7857<br>($\pm 0.0116$) |
| CNN-Dense (S+F) | | 0.4201<br>($\pm 0.0069$) | 0.7155<br>($\pm 0.0095$) |
| TCN-Dense (S+F) | | 0.3821<br>($\pm 0.0144$) | 0.6287<br>($\pm 0.0157$) |
| XGBoost (F) | F1-score | 0.5371<br>($\pm 0.0056$) | 0.9109<br>($\pm 0.0068$) |
| XGBoost (S+F) | | 0.5255<br>($\pm 0.0069$) | 0.9044<br>($\pm 0.0063$) |
| CNN-Dense (C+F) | | 0.5517<br>($\pm 0.0146$) | 0.7845<br>($\pm 0.0108$) |
| CNN-Dense (S+F) | | 0.4265<br>($\pm 0.0071$) | 0.7112<br>($\pm 0.0071$) |
| TCN-Dense (S+F) | | 0.3948<br>($\pm 0.0157$) | 0.6296<br>($\pm 0.0166$) |

<sup>1</sup>Performance based on test set evaluation metrics of different ML/DL classification models at different minimum event durations. Values are means and std. dev. (N=5).
<sup>2</sup>Models are named as defined in the text.
<sup>3</sup>Metric names are abbreviated and refer to overall accuracy and macro-averaged F1-score.

The TCN-Dense (S+F) model, a capable architecture previously developed for event-level classification of simulated peptides (29), served as a baseline in our evaluation. This model, trained on translocation event state sequences and corresponding event-level features, showed a modest average accuracy (0.6287 (±0.0157) at 20 ms cutoff) and generally followed the trend of improved predictions with increasing event length **(Table 1)**. The CNN-Dense (S+F) model, likewise trained on state sequences and event-level features, surprisingly achieved slightly better performance (e.g., 0.7155 (±0.0095) accuracy at 20 ms cutoff, N=5), while being less computationally demanding **(Table 1)**. Critically, the CNN-Dense (C+F) model, which was trained directly on the scaled current sequences and event-level features, emerged as the most robust DL model. Its superior performance (e.g., 0.7857 (±0.0116) accuracy at 20 ms cutoff, N=5) strongly suggests that the raw current sequences of translocation events possess more discriminative information than the more abstract conductance state sequences **(Table 1)**.

Detailed confusion matrices for the CNN-Dense (C+F) model at both 5 ms and 20 ms minimum event durations **(Fig. 3B,C)** further elucidate class-by-class prediction quality. These reveal that the aromatic guest-host peptides (Phe, Trp, TrpDL, and Tyr) were consistently predicted with high fidelity, even considering their nuanced chemical differences. Notably, the models demonstrated the nanopore’s ability to discriminate between guest-host Trp and guest-host TrpDL, which differ only in backbone stereochemistry. This finding strongly suggests that peptide backbone conformational dynamics, previously reported as key biophysical properties detectable by the PA nanopore (18, 19), are indeed unique features leveraged by the classifier.

Conversely, consistent misclassification was observed between guest-host Ala and guest-host Thr. Their rapid translocation events exhibited similar dynamics and consequently occupied overlapping feature space, reflecting the subtle chemical differences between these two residues. For such cases, the analysis consistently indicated that increasing the minimum event duration parameter, thereby focusing the classification models on longer, more information-rich events, represents the best strategy for improving classification accuracy for these challenging cases.

### Performance of ML and ML/DL-hybrid classifiers

Beyond DL approaches, we investigated the performance of tree-based machine learning classifiers. Specifically, XGBoost, a highly efficient ensemble learning method leveraging gradient-boosted decision trees (30), was employed using only the pre-extracted event-level feature set—referred to as XGBoost (F). Like the DL models, a preliminary single-replicate scan across the same range of minimum event durations (5-20 ms) revealed a consistent trend: classification metrics significantly improved as shorter, lower-information events were progressively removed from the training data **(Table S3)**. At a minimum event duration of 20 ms, the XGBoost (F) model achieved exceptionally high performance, with an average overall accuracy of 0.9112 (±0.0069) (N=5) **(Table 1)**. A detailed normalized confusion matrix for this configuration **(Fig. 4)** illustrates the robust per-class prediction quality. Consistent with observations from other models, guest-host Ala and guest-host Thr remained the weakest performing classes, exhibiting some low-level confusion **(Fig. 4)**. However, their F1-scores were still notably high, exceeding 0.8. A significant practical advantage of XGBoost over the DL-based approaches was its substantial computational efficiency during both model training and inference.

**Fig. 4.**
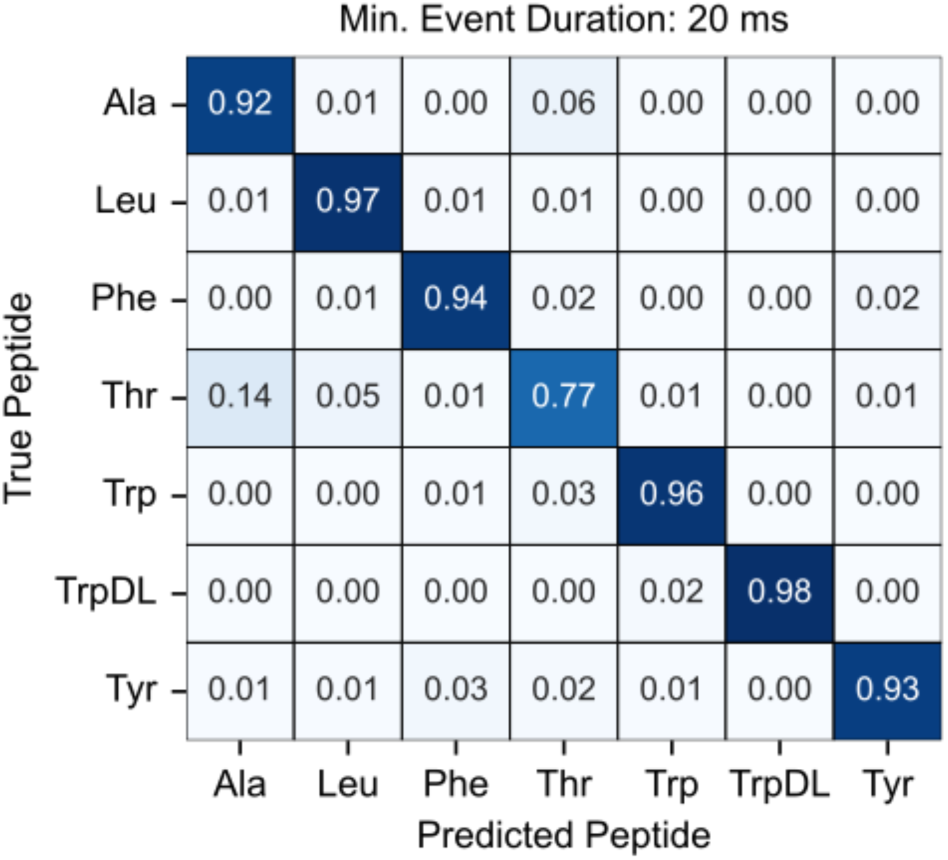
High-performance tree-based classification of translocation events. Normalized confusion matrix for XGBoost classification using the event-level feature set. This matrix represents the best-performing XGBoost (F) model at a minimum event duration of 20 ms. Rows represent true peptide labels, and columns represent predicted labels. Values indicate the proportion of events from a given true class that were predicted as each class. Despite the model’s overall high performance, a very moderate level of misclassification is observed between the chemically similar guest-host Ala and guest-host Thr classes.

We also explored a hybrid ML/DL model architecture to assess if sequence-derived embeddings could enhance feature-based classification by XGBoost. This hybrid approach XGBoost (S+F) utilized a CNN to generate embeddings from the state sequences, which were then merged with the event-level feature set, and this combined vector was subsequently used to train an XGBoost classifier. While this hybrid model also demonstrated increasing performance with longer minimum event durations, it did not outperform the XGBoost (F) model acting solely on the event-level feature set **(Table 1)**. This indicates that, for our nanopore/peptide system, the addition of CNN-derived embeddings from state sequences did not provide significant additional discriminative information when combined with the already comprehensive engineered event-level features. In conclusion, for our nanopore and guest-host peptide system, the combination of carefully engineered features (particularly those derived from longer events) with a robust tree-based model like XGBoost (F) achieved superior or highly comparable classification performance with significantly greater computational efficiency than the tested deep learning sequence models.

## Discussion

### High-performance nanopore biosensing

Towards addressing the unmet challenge of rapid/accurate biomolecule characterization, we have described the utility of anthrax toxin’s PA nanopore as powerful peptide biosensor platform. A robust machine learning framework was developed to this end, which could classify seven diverse guest-host peptides based on individual nanopore translocation events, achieving ∼0.90 accuracy **(Table 1)**. The key breakthrough is having the capability of classifying individual events, instead of relying on global features or ensemble analyses, paving the way for classifying complex peptide mixtures.

Carefully engineered, event-level biophysical features proved highly discriminative for peptide classification. The XGBoost (F) model, relying solely on these event-level features, achieved the highest classification accuracy, outperforming even complex deep learning architectures, which trained on state or current sequences and features. This underscores the power of expert-derived biophysical features for this specific application. Even when XGBoost (S+F) was trained on features plus state sequence embeddings, its predictive capacity did not exceed the XGBoost (F) trained on features alone. This result establishes that all the discriminative information resides in the feature set itself. Another key insight from this study was the observation that the systematic filtering of shorter events (i.e., by increasing the minimum event duration) significantly improved UMAP clustering **(Fig. 2)** and ML/DL classification performance across all tested models **(Table 1, Table S3)**. This indicates that longer translocation events carry more stable and discriminative information, likely due to more complete and resolved multi-state interactions with various peptide-clamp sites the pore. Taken together these observations demonstrate the viability of high-performance peptide classification using this nanopore system.

### Exquisite sensitivity and broad discriminatory power of the PA nanopore

The models consistently demonstrated the ability to discriminate between peptides differing only in backbone stereochemistry (e.g., Trp and TrpDL), highlighting the remarkable intrinsic sensitivity of the PA nanopore to subtle conformational and interaction dynamics **(Figs. 3C, 4)**. Previous biophysical characterizations of the PA nanopore revealed altered peptide and clamp dynamics (18) and helix-coil transition dynamics (19), when only stereochemistry was manipulated. This conformation biosensing capability is likely somewhat unique to this nanopore system. Furthermore, the discriminatory power extended to other chemically subtle differences between peptides. Guest-host Phe and guest-host Tyr were well classified despite differing only by a hydroxyl group. Guest-host Phe and guest-host Leu, despite having similar hydrophobicity, were distinguishable by this pore and computational approach. Early work on model compound binding to ϕ-clamp site (F427) showed enhanced binding of aromatic compounds, suggesting ring-ring interactions are made to the phenyl groups (14). Overall, these observations underscore the nanopore’s ability to discern fine chemical, structural, and conformational nuances beyond what might be expected.

### Comparative performance of ML versus DL

In this study, XGBoost (F), which trained only on the engineered features, outperformed all other sequence-based DL and hybrid models. These results may reveal that the engineered event-level features are highly optimized and capture the most relevant biophysical information more directly. The current DL sequence models might be too general, or the raw sequence data might be too noisy/lack sufficient resolved signal for subtle distinctions compared to preprocessed features from this multi-state system. Nonetheless, while neural networks underperformed in this classification task using a fixed feature set, their strength lies in potential for better adaptability. They will likely be pivotal in classifying more diverse, real-world peptides that exhibit variable numbers of observed conductance states (e.g., 2-state vs. 3-state vs. 4-state or higher). This is a current limitation of a fixed feature set, which here was specifically for 4-state peptides. DL models will offer broader applicability for handling heterogeneous data where a universal fixed feature set is not feasible. For specific, well-characterized systems with robust feature engineering, traditional ML can be superior. However, for broader, more complex, or less understood systems, a DL model that can learn directly from raw, variable-length, and variable-state data will be essential.

Comparatively, the CNN-Dense (C+F) model was superior to the TCN-Dense (S+F) model in terms of accuracy and computational efficiency. We anticipate the former architecture will be viable for more general classification applications—even classifying from current sequences alone.

### Challenges and future opportunities

The most challenging pair of peptides to classify were guest-host Ala and guest-host Thr in all tested models. The current best strategy to reduce their misclassification and confusion is to increase the minimum event duration to filter out lower quality short events. The wild-type PA nanopore used here may not be perfectly suited to distinguish all chemistries. Therefore, a multiplexed approach can be advantageous, where a variant, orthogonal PA nanopore (e.g., a ϕ-clamp mutant, like F427A) is used (31). This approach may offer better discrimination between these two challenging peptides. Having an alternate nanopore may offer orthogonal discriminatory information to augment inference. While this study focused on classification, insights gained here (feature engineering, event-length optimization, and analyte sensitivity) lay the groundwork for more ambitious goals like direct peptide sequencing. Neural networks are likely essential for this task. For example, a sequence-to-sequence (current-to-peptide sequence) architecture could be a powerful avenue to pursue. Beyond refining classification for known peptides, future work will also explore the generalizability of our approach to wider, chemically diverse peptide sets, complex mixtures, and even different nanopore architectures. Furthermore, the integration of more advanced feature extraction techniques, potentially leveraging deep learning for automated feature engineering or multi-modal data fusion, could further unlock unprecedented performance beyond our current methods. Finally, translating these advancements to real-world applications will necessitate addressing practical challenges inherent in complex biological matrices, such as sample preparation, signal-to-noise optimization, and high-throughput integration. Ultimately, by persistently refining both the nanopore biosensor and the computational intelligence that deciphers its signals, we move closer to a future where rapid, single-molecule proteomic analysis becomes a routine and transformative reality.

## Materials and Methods

### Nanopore and peptides

Monomeric 83-kDa PA (PA_83_) preprotein and the oligomerized heptameric form (PA_7_) were prepared as described (32). PA_83_ monomer was overexpressed in *Escherichia coli* BL21(DE3), using a pET22b plasmid, which directs expression to the periplasm. Cell cultures were grown at 37 °C in a custom 5 L fermentor using ECPM1 growth media (33), which was supplemented with carbenicillin (50 mg/L). Once reaching an OD_600_ of 3-10, the cultures were then induced with 1 mM isopropyl β-d-thiogalactopyranoside for ∼3 h at 30 °C. PA_83_ was released from the periplasm by resuspending pelleted cells on ice using a wire whisk with 1 L of hypertonic sucrose buffer (20% sucrose, 20 mM Tris-Cl, 0.5 mM EDTA, pH 8) followed by osmotic shock of centrifuged/pelleted cells using a wire whisk in 1 L of hypotonic solution (5 mM MgCl_2_). Released PA_83_, isolated after centrifugation to remove cellular debris, was purified on Q-Sepharose anion-exchange chromatography in 20 mM Tris-Cl, pH 8.0 by binding and then eluting with a linear salt gradient using 20 mM Tris-Cl, pH 8.0 with 1 M NaCl.

To make PA_7_ prepore oligomers, purified PA_83_ at a concentration of 1 mg/ml was treated with trypsin (1:1000 wt/wt trypsin:PA) for 30 min at room temperature to form nicked PA. Trypsin was subsequently inhibited with soybean trypsin inhibitor at 1:100 dilution (wt/wt soybean trypsin inhibitor:PA). Nicked PA was applied to Q-Sepharose to then isolate the PA_7_; oligomer was bound to the column in 20 mM Tris-chloride, pH 8.0 and eluted by a linear salt gradient using 20 mM Tris-Cl, 1 M NaCl, pH 8.0. PA_7_ was concentrated and frozen in small aliquots to maintain reproducible nanopore insertion activity in planar bilayer experiments.

Ten-residue guest-host peptides of the general sequence, KKKKKXXSXX, where X = A, L, F, T, W, and Y, were synthesized with standard ʟ amino acids as indicated (18, 21) (Elim Biopharmaceuticals). One stereochemical variant of X = W (abbreviated TrpDL) was produced, where instead of synthesizing the peptide with uniform ʟ amino acids, an alternating pattern of ᴅ and ʟ amino acids was used (18).

### Single-channel electrophysiology

Planar lipid bilayer currents were recorded using an Axopatch 200B amplifier interfaced by a Digidata 1440A acquisition system (Molecular Devices) (18, 32, 34). Membranes were formed by painting across a 50-μm aperture of a 1-mL white Delrin cup with 3% (wt/vol) 1,2-diphytanoyl-*sn*-glycero-3-phosphocholine (Avanti Polar Lipids) in *n*-decane. The *cis* (side to which the PA_7_ is added) and *trans* chambers were bathed in symmetric single-channel buffer (SCB: 100 mM KCl, 1 mM EDTA, 10 mM succinic acid, pH 5.60). Recordings were acquired at 400-600 Hz using PCLAMP10. The applied voltage is defined as Δψ = ψ_cis_ - ψ_trans_ (where ψ_trans_ was set to 0 mV).

Single-channel recordings of the guest-host peptide translocations via the PA nanopore were carried out as described (18) with some slight differences. A single PA channel was inserted into a painted bilayer at a Δψ of 20-30 mV by adding ∼2 pM of PA_7_ (freshly diluted from a 2-μM stock) to the *cis* side of the membrane. The oligomer converts to the nanopore state by inserting into the membrane in an oriented manner. Once a single channel inserted, the *cis* chamber was perfused by fresh SCB to remove excess uninserted PA_7_. Then the desired peptide analyte was added to the *cis* chamber at 5 to 20 nM. Translocation data were acquired by stepping the applied Δψ to a higher positive value and collecting recordings of the translocation event stream for up to several minutes.

Minor processing as well as conductance state labeling of the raw single-channel event stream recordings was subsequently performed. Rare transient out-of-range current spikes, insertion of second channels, and inactivated channels were removed by a ‘force values’ routine in CLAMPFIT. Some translocation recordings that were acquired at 500 and 600 Hz were downsampled to 400 Hz by decimation in Python using the scipy.signal library. 400 Hz was chosen to maximize the data volume at a consistent time step for ML/DL-based peptide classification. Four discrete conductance states were then detected in these recordings using single-channel event detection in CLAMPFIT. We noted that during event detection the shortest translocations consisting of a single time step (2.5 ms at 400 Hz) were identified by CLAMPFIT as being two time point long events (5 ms). By convention during event detection in CLAMPFIT, the fully blocked peptide-bound state was state 0, the intermediate closest to the fully blocked state was state 1, the intermediate closest to the open state was state 2, and the open state was state 3. Start and stop times of all detected events were used to label the state of each time point in the raw current recordings, producing a three-column CSV file of the stream with columns labeled as ‘time’, ‘current’, and ‘state’. All labeled CSV stream files for the seven peptides were entered into a local annotated peptide database to aid in *in situ* loading/preprocessing for each tested ML/DL model (described below).

### Hardware and software used for ML/DL

Anaconda was used to create a Python 3.10.16 environment, where TensorFlow (2.16.2) (35), Keras TCN (3.5.6) (36), XGBoost (3.0.0) (30), and other standard modules were installed. The hardware used in training ML/DL peptide classification models was a 2025 MacBook Pro with M4 Apple Silicon and 24 GB of RAM. GPU cores were utilized during training by installing tensorflow-metal. All source code is available at GitHub (https://github.com/bakrantz/Pept-Class).

### Preprocessing, translocation event segmentation, and feature extraction

Raw translocation event streams were preprocessed prior to segmentation and feature extraction **(Fig. S1B)**. While the segmentation core can apply low- and high-pass filtering and baseline correction, these filters/corrections were not applied to the experimental guest-host peptide translocation datasets presented in this study. The most critical preprocessing parameter was the minimum event duration, which served as an effective filter for excluding very short-duration events. This parameter was systematically varied in the range of 5 to 20 ms to assess its empirical impact on downstream machine learning and deep learning classification performance.

Following the application of these processing parameters, the state-labeled raw event streams were segmented into individual translocation events. Each event was defined as initiating when the current changed from the fully open state (state 3, corresponding to baseline current) to any peptide-bound state (state 0, 1, or 2) and terminating when the current returned to the open state. From these segmented events, both raw current sequences and corresponding state sequences were extracted.

A comprehensive set of event-level features was then computed from these sequences using a custom segmentation core. This core maintains a generalizable framework to process peptide translocation events from systems exhibiting diverse mechanisms and an arbitrary number of states. These features were initially categorized into scalar, vector, and matrix data structures, with values in vector and matrix features being state or transition enumerated. Scalar features included: Shannon entropy of state sequence, event duration, number of transitions, time of the first transition, and total number of states visited during the event. Vector features included: observed conductance state Boolean, observed conductance levels, probability of residing in each state, and longest dwell time in each state. Matrix features included: average dwell time for specific state-to-state transitions, variance of dwell time for transitions, and ratio of probabilities between states.

While the segmentation core also supports the extraction of global features that have yielded high-quality classification results in simulated datasets (e.g., >0.99 accuracy) (29), these were not utilized in the classifications of the experimental guest-host datasets presented here, as our focus was solely on the more challenging and practical task of individual event-level classification.

For downstream classification, all matrix features were flattened into one-dimensional arrays and appended with the vector and scalar features to form a single feature vector for each translocation event. These flattened descriptive key names for the features were generated to maintain traceability in subsequent applications. All processed event sequences, their flattened features, and associated feature key names were saved as a Python pickle object for efficient storage and retrieval. A local peptide events database was employed to track these preprocessed pickle files, thereby preventing redundant segmentations and feature extractions from raw datasets.

### DL-based peptide classification

Peptide classification using DL models followed a common architectural schema: a sequence (either conductance states or raw current) was processed by a CNN or a TCN branch, the output of which was then concatenated with concurrently input event-level features **(Fig. 3A, Fig. S1C)**. This combined representation was subsequently fed into a fully connected (Dense) layer, culminating in a final softmax output for multiclass classification.

Three distinct DL models were evaluated. (i) The TCN-Dense (S+F) model was trained on the conductance state sequences (S) combined with event-level features (F). The architecture and hyperparameters for this model were identical to those previously described (29), with modifications only to the data loading from our current database. (ii) The CNN-Dense (S+F) model also utilized conductance state sequences (S) and event-level features (F). The sequence processing branch consisted of three one-dimensional CNN layers, each followed by a dropout layer with a rate of 0.3. The output of these CNN layers was then subjected to global max pooling before concatenation with the event-level features. The concatenated output was then passed to the final Dense classification layers. (iii) The CNN-Dense (C+F) model was structurally similar to the CNN-Dense (S+F) model but was trained on current sequences (C) instead of state sequences, combined with event-level features (F). Its sequence processing branch also employed three one-dimensional CNN layers with dropouts of 0.3 and global max pooling prior to concatenation with the event-level features, followed by analogous Dense classification layers.

For all three models, data preparation and training procedures were standardized as follows. Data Balancing: during data loading, class balancing was performed *in situ* by downsampling all classes to match the number of events in the class with the fewest samples **(Table S2)**. Data Split: the balanced dataset was partitioned into an 80% training set and a 20% testing set. Feature Preprocessing: event-level features were scaled using a StandardScaler.

Missing (NaN) values within the feature sets were imputed with a value of −1. Model Optimization: the Adam optimizer was used for model training, with categorical cross-entropy as the loss function. Model performance was monitored using metrics including accuracy, precision, and recall. Training Protocol: training was performed for approximately 100 epochs with a batch size of 32. To prevent overfitting and optimize training, several callbacks were implemented: early stopping (to halt training when validation performance no longer improved), learning rate reduction on plateau (to reduce the learning rate if validation loss stalled), and model checkpointing (to save the best performing model weights).

The classification performance of each trained model was subsequently evaluated on its respective unseen testing dataset. Comprehensive metrics, including overall accuracy, precision, recall, and F1-score for each peptide class, were summarized in a standard classification report. Additionally, confusion matrices were generated to provide a detailed visualization of per-class prediction accuracy and misclassification patterns.

### ML-based peptide classification

ML-based classification of peptide translocation events was performed using the gradient boosting framework, XGBoost (30). For this, peptide event data, previously extracted and characterized into event-level features, were loaded from a local database. To ensure balanced class representation, all peptide classes were downsampled to match the class with the minimum number of events **(Table S2)**. The comprehensive dataset was then split into an 80% training set and a 20% testing set for model development and evaluation, respectively. For clarity, throughout this features-only model is referred to as XGBoost (F). The XGBoost classifier was configured with the following key parameters: a multiclass classification objective, the number of target classes set to the total number of peptides, 1000 boosting rounds (trees), and a learning rate of 0.05 to control the step size shrinkage. Regularization was applied with max depth of 5 to limit tree complexity, minimum child weight was 1 to control minimum sum of instance weight (hessian) needed in a child, gamma was set to 0 for minimum loss reduction required to make a further partition on a leaf node, subsample was 0.8 (fraction of samples used per tree), and the fraction of features used per tree was 0.8. L1 and L2 regularization were used. For reproducibility, a random state was fixed, and computation was distributed across all available CPU cores. The model’s performance during training was monitored using the multiclass classification error. The trained model’s performance was evaluated on the unseen testing dataset. Classification metrics including accuracy, precision, recall, and F1-score were summarized in a standard classification report, and a confusion matrix was generated to visualize per-class prediction accuracy.

### Hybrid DL/ML-based peptide classification

A hybrid classification approach, XGBoost (S+F), was developed to leverage the representational power of deep learning for sequence data alongside the robust performance of tree-based ensemble methods. This model combined sequence-derived embeddings from a supervised CNN with the previously defined event-level features, which were then merged and fed into the XGBoost classifier.

#### CNN-based sequence embedding generation

To generate sequence embeddings, a dedicated CNN model was trained to perform multiclass peptide classification directly on conductance state sequences. Raw state sequences were first remapped to integer IDs, where original states (e.g., 0, 1, 2, 3) were shifted by one (e.g., 1, 2, 3, 4) to reserve ‘0’ as a dedicated padding token. These remapped sequences were then post padded to a fixed sequence length of 1300 time points (which contained 99% of the events), thus ensuring a uniform input dimension for the CNN. During data loading for CNN training, all peptide classes were downsampled to the size of the smallest class to maintain class balance. The dataset was subsequently split into an 80% training set and a 20% validation set.

The CNN embedding model architecture consisted of an embedding layer (input dimension vocab size was number of unique remapped states + 1 (to include padding token) and output dimension was 128) that also masked the padding token (value 0). This was followed by a stack of two 1D CNN layers, each with 128 filters and a kernel size of 3, utilizing rectified linear unit activation and ‘same’ padding. Each layer’s output was subjected to batch normalization and max pooling with a pool size of 2, followed by a dropout layer of 0.4. A global max pooling layer then summarized the processed sequence into a fixed-size representation.

This was fed into a Dense layer with an output embedding dimension of 128, producing the final sequence embedding. A separate Dense classification head (with softmax activation) was attached to this embedding layer, allowing the entire CNN to be trained in a supervised manner for peptide classification. The model was compiled using the Adam optimizer with sparse categorical cross-entropy for the classification head (the embedding output had loss of none as it was not directly optimized during this phase). Training was performed for 100 epochs with a batch size of 32, incorporating early stopping (restoring best weights) and reduce learning rate on plateau callbacks to prevent overfitting and optimize learning. The training history (loss and accuracy) were monitored on the validation set. After training, the final sequence embeddings were extracted from the trained model’s embedding layer for both the training and testing sets.

#### Hybrid model training and evaluation

For the hybrid model, event data (including sequences and features) for all peptides were loaded from a local database, applying the same minimum event duration filter range as used for the individual DL and ML models (e.g., ≥ 20 ms was most optimal empirically). The dataset was then split 80-20 into training and testing sets.

The extracted CNN sequence embeddings (from the previously trained encoder) were then horizontally concatenated with their corresponding event-level features for both the training and testing sets. This combined feature vector served as the input for the final XGBoost classifier. The XGBoost classifier was configured with identical parameters to those used when trained solely on event-level features (as detailed in the previous section). Early stopping was also applied during XGBoost training. The performance of the hybrid XGBoost (S+F) model was assessed on the independent test set using a comprehensive classification report, providing overall accuracy, precision, recall, and F1-score. A confusion matrix was also generated to visualize per-class prediction accuracy.

## Acknowledgments

We thank members of the department as well as Tobin Sosnick for useful feedback and discussions. J.M.C. and B.A.K. conceived of the experiments. J.M.C. collected the data. B.A.K performed the analysis. B.A.K. and J.M.C. wrote the manuscript. Portions of this document, including some of the Python code and language refinement, were generated with the assistance of AI-powered tools. All content was reviewed and approved by the authors, who take full responsibility for its accuracy.

---

**Fig. S1.**
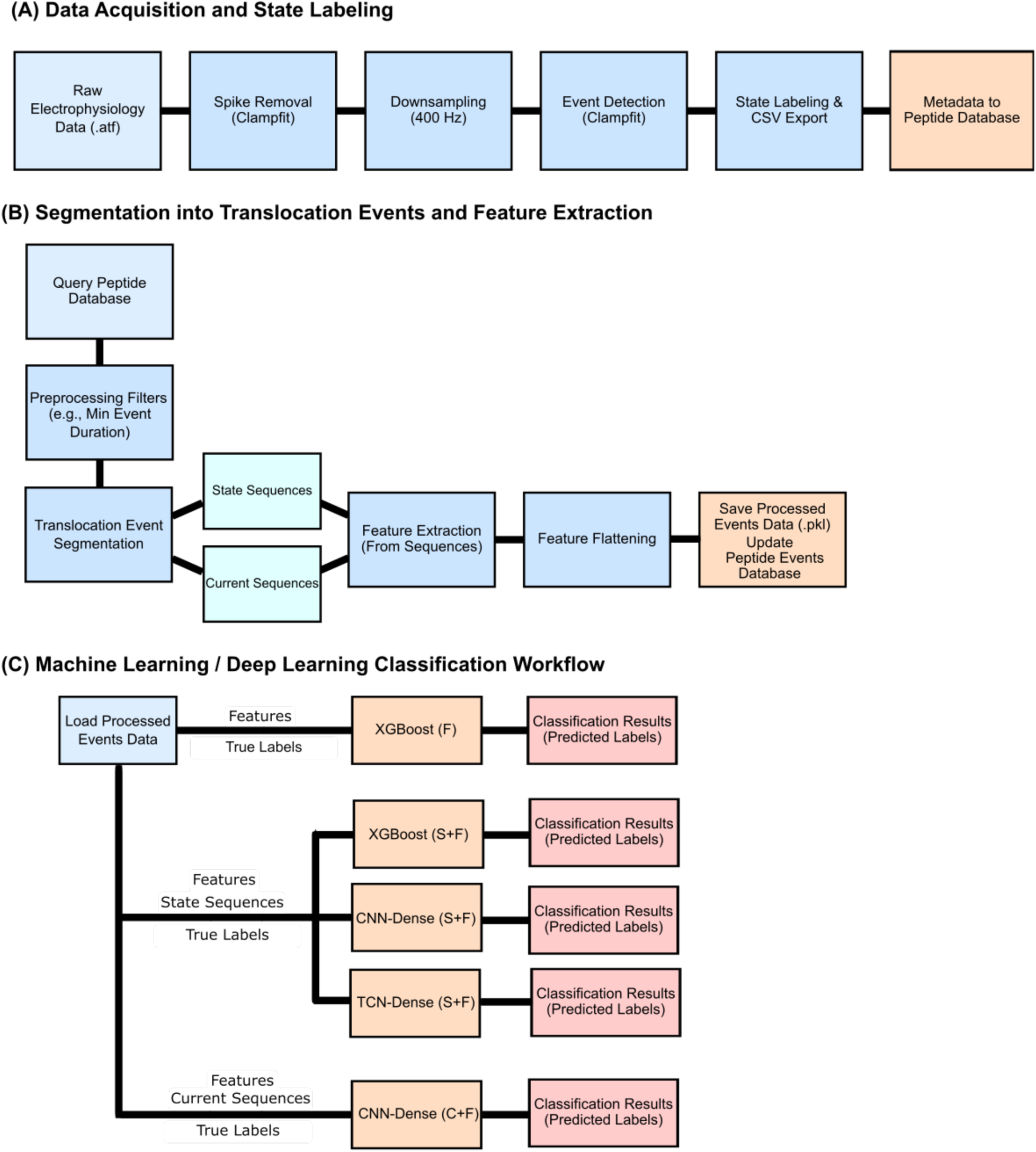
Processing and analysis schemes. **(A)** Data acquisition and state labeling. **(B)** Event segmentation and feature calculation. **(C)** ML/DL classification workflow.

**Table S1.** Recording time^1^ per peptide class in the dataset.

| Guest-Host Peptide | Time (seconds) | Time (hours) |
| --- | --- | --- |
| Ala | 1717.8125 | 0.4772 |
| Leu | 513.3825 | 0.1426 |
| Phe | 514.57 | 0.1429 |
| Thr | 5141.785 | 1.4283 |
| Trp | 1929.47 | 0.536 |
| TrpDL | 1975.4625 | 0.5487 |
| Tyr | 2559.91 | 0.7111 |
| TOTAL Dataset | 14352.3925 | 3.9868 |

<sup>1</sup>These recording times include all data at the 70 mV voltage condition and peptide concentration range (5-20 nM).

**Table S2.** Translocation event counts at different minimum event duration filters.

| Guest-Host Peptide | 5 ms Minimum Event Duration |  | 20 ms Minimum Event Duration |  |
| --- | --- | --- | --- | --- |
|  | Events Before Downsampling | Events After Downsampling<sup>1</sup> | Events Before Downsampling | Events After Downsampling<sup>2</sup> |
| Ala | 83,546 | 2,435 | 6,164 | 1,340 |
| Leu | 15,983 | 2,435 | 2,231 | 1,340 |
| Phe | 5,307 | 2,435 | 2,342 | 1,340 |
| Thr | 368,562 | 2,435 | 31,546 | 1,340 |
| Trp | 10,483 | 2,435 | 5,109 | 1,340 |
| TrpDL | 2,435 | 2,435 | 1,340 | 1,340 |
| Tyr | 77,338 | 2,435 | 37,978 | 1,340 |
| Total | 563,654 | 17,045 | 86,710 | 9,380 |

<sup>1</sup>For the 5 ms minimum event duration dataset, all peptide classes were downsampled to 2,435 events, based on the class with the fewest events (TrpDL), to create a balanced dataset for model training and evaluation.
<sup>2</sup>For the 20 ms minimum event duration dataset, all peptide classes were downsampled to 1,340 events, based on the class with the fewest events (TrpDL), to create a balanced dataset for model training and evaluation.

**Table S3.** Model performance metrics at varying minimum event duration.^1^.

| Model<sup>2</sup> | Metric<sup>3</sup> | Minimum event duration |  |  |  |  |  |
| --- | --- | --- | --- | --- | --- | --- | --- |
|  |  | 5 ms | 7.5 ms | 10 ms | 12.5 ms | 15 ms | 20 ms |
| XGBoost (F) | Accuracy | 0.5483 | 0.7934 | 0.8235 | 0.8697 | 0.8737 | 0.9083 |
| XGBoost (S+F) |  | 0.5345 | 0.7905 | 0.8084 | 0.8755 | 0.8536 | 0.9078 |
| CNN-Dense (C+F) |  | 0.5456 | 0.6949 | 0.7312 | 0.7564 | 0.769 | 0.8006 |
| CNN-Dense (S+F) |  | 0.4303 | 0.6397 | 0.6471 | 0.686 | 0.6848 | 0.7281 |
| TCN-Dense (S+F) |  | 0.3711 | 0.5407 | 0.568 | 0.6242 | 0.6182 | 0.6578 |
| XGBoost (F) | Precision | 0.5526 | 0.797 | 0.8262 | 0.8692 | 0.8739 | 0.9091 |
| XGBoost (S+F) |  | 0.5531 | 0.7913 | 0.8121 | 0.8746 | 0.8548 | 0.9089 |
| CNN-Dense (C+F) |  | 0.5602 | 0.7118 | 0.7457 | 0.7635 | 0.776 | 0.802 |
| CNN-Dense (S+F) |  | 0.4982 | 0.6455 | 0.6615 | 0.6968 | 0.6897 | 0.7412 |
| TCN-Dense (S+F) |  | 0.5103 | 0.6088 | 0.6056 | 0.6576 | 0.6627 | 0.692 |
| XGBoost (F) | Recall | 0.5483 | 0.7934 | 0.8234 | 0.8697 | 0.8737 | 0.9083 |
| XGBoost (S+F) |  | 0.5345 | 0.7905 | 0.8083 | 0.8755 | 0.8536 | 0.9078 |
| CNN-Dense (C+F) |  | 0.5456 | 0.6949 | 0.731 | 0.7563 | 0.7689 | 0.8006 |
| CNN-Dense (S+F) |  | 0.4303 | 0.6396 | 0.647 | 0.6858 | 0.6847 | 0.7281 |
| TCN-Dense (S+F) |  | 0.3711 | 0.5407 | 0.568 | 0.6241 | 0.6181 | 0.6578 |
| XGBoost (F) | F1-score | 0.5436 | 0.7941 | 0.8232 | 0.8687 | 0.8731 | 0.9085 |
| XGBoost (S+F) |  | 0.5159 | 0.7899 | 0.8083 | 0.8742 | 0.8533 | 0.9082 |
| CNN-Dense (C+F) |  | 0.5457 | 0.6983 | 0.7364 | 0.7583 | 0.7687 | 0.7981 |
| CNN-Dense (S+F) |  | 0.4308 | 0.6353 | 0.6483 | 0.6857 | 0.6825 | 0.7233 |
| TCN-Dense (S+F) |  | 0.3875 | 0.5596 | 0.5765 | 0.6318 | 0.6264 | 0.6597 |

<sup>1</sup>Performance based on test set evaluation metrics of different ML/DL classification models at different minimum event durations (N=1). Finalized replicated performance metrics (N=5) for these models at either minimum event duration extreme are presented in **Table 1**.
<sup>2</sup>Models are named as defined in the text.
<sup>3</sup>Metric names are abbreviated and refer to overall accuracy, macro-averaged precision, macro-averaged recall, and macro-averaged F1-score.

## References

1. V. Castiglione et al., Biomarkers for the diagnosis and management of heart failure. Heart Fail Rev 27, 625–643 (2022).

2. P. Povoa et al., How to use biomarkers of infection or sepsis at the bedside: guide to clinicians. Intensive Care Med 49, 142–153 (2023).

3. R. Mastrantonio, H. You, L. Tamagnone, Semaphorins as emerging clinical biomarkers and therapeutic targets in cancer. Theranostics 11, 3262–3277 (2021).

4. L. Ratinho, N. Meyer, S. Greive, B. Cressiot, J. Pelta, Nanopore sensing of protein and peptide conformation for point-of-care applications. Nat Commun 16, 3211 (2025).

5. B. Lin, J. Hui, H. Mao, Nanopore Technology and Its Applications in Gene Sequencing. Biosensors (Basel) 11 (2021).

6. Y. Goto, R. Akahori, I. Yanagi, Challenges of Single-Molecule DNA Sequencing with Solid-State Nanopores. Adv Exp Med Biol 1129, 131–142 (2019).

7. X. Wei et al., Engineering Biological Nanopore Approaches toward Protein Sequencing. ACS Nano 17, 16369–16395 (2023).

8. J. Jiang, B. L. Pentelute, R. J. Collier, Z. H. Zhou, Atomic structure of anthrax protective antigen pore elucidates toxin translocation. Nature 521, 545–549 (2015).

9. N. J. Hardenbrook et al., Atomic structures of anthrax toxin protective antigen channels bound to partially unfolded lethal and edema factors. Nat Commun 11, 840 (2020).

10. K. Zhou et al., Atomic Structures of Anthrax Prechannel Bound with Full-Length Lethal and Edema Factors. Structure 28, 879–887 e873 (2020).

11. A. J. Machen, M. T. Fisher, B. D. Freudenthal, Anthrax toxin translocation complex reveals insight into the lethal factor unfolding and refolding mechanism. Sci Rep 11, 13038 (2021).

12. G. K. Feld et al., Structural basis for the unfolding of anthrax lethal factor by protective antigen oligomers. Nature Struct. Mol. Biol. 17, 1383–1390 (2010).

13. S. L. Wynia-Smith, M. J. Brown, G. Chirichella, G. Kemalyan, B. A. Krantz, Electrostatic ratchet in the protective antigen channel promotes anthrax toxin translocation. J Biol Chem 287, 43753–43764 (2012).

14. B. A. Krantz et al., A phenylalanine clamp catalyzes protein translocation through the anthrax toxin pore. Science 309, 777–781 (2005).

15. B. A. Krantz, Anthrax Toxin: Model System for Studying Protein Translocation. J Mol Biol 436, 168521 (2024).

16. S. Zhang, E. Udho, Z. Wu, R. J. Collier, A. Finkelstein, Protein translocation through anthrax toxin channels formed in planar lipid bilayers. Biophys. J. 87, 3842–3849 (2004).

17. B. A. Krantz, A. Finkelstein, R. J. Collier, Protein translocation through the anthrax toxin transmembrane pore is driven by a proton gradient. J. Mol. Biol. 355, 968–979 (2006).

18. K. Ghosal et al., Dynamic Phenylalanine Clamp Interactions Define Single-Channel Polypeptide Translocation through the Anthrax Toxin Protective Antigen Channel. J Mol Biol 429, 900–910 (2017).

19. D. Das, B. A. Krantz, Peptide- and proton-driven allosteric clamps catalyze anthrax toxin translocation across membranes. Proc Natl Acad Sci U S A 113, 9611–9616 (2016).

20. S. R. Blanke, J. C. Milne, E. L. Benson, R. J. Collier, Fused polycationic peptide mediates delivery of diphtheria toxin A chain to the cytosol in the presence of anthrax protective antigen. Proc. Natl Acad. Sci. U.S.A. 93, 8437–8442 (1996).

21. J. M. Colby, B. A. Krantz, Peptide Probes Reveal a Hydrophobic Steric Ratchet in the Anthrax Toxin Protective Antigen Translocase. J Mol Biol 427, 3598–3606 (2015).

22. D. Das, B. A. Krantz, Secondary Structure Preferences of the Anthrax Toxin Protective Antigen Translocase. J Mol Biol 429, 753–762 (2017).

23. N. Celik et al., Deep-Channel uses deep neural networks to detect single-molecule events from patch-clamp data. Commun Biol 3, 3 (2020).

24. S. Yang et al., Deep Learning-Based Ion Channel Kinetics Analysis for Automated Patch Clamp Recording. Adv Sci (Weinh) 12, e2404166 (2025).

25. D. Dematties, C. Wen, M. D. Perez, D. Zhou, S. L. Zhang, Deep Learning of Nanopore Sensing Signals Using a Bi-Path Network. ACS Nano 15, 14419–14429 (2021).

26. C. Cao et al., Aerolysin nanopores decode digital information stored in tailored macromolecular analytes. Sci Adv 6 (2020).

27. D. Rodriguez-Larrea, Single-aminoacid discrimination in proteins with homogeneous nanopore sensors and neural networks. Biosens Bioelectron 180, 113108 (2021).

28. G. Yamini et al., Hydrophobic Gating and 1/f Noise of the Anthrax Toxin Channel. J Phys Chem B 125, 5466–5478 (2021).

29. B. A. Krantz, Deep Learning-Based Classification of Peptide Analytes from Single-Channel Nanopore Translocation Events. bioRxiv 10.1101/2025.05.04.652126 (2025).

30. T. Chen, C. Guestrin (2016) XGBoost: A Scalable Tree Boosting System. in Proceedings of the 22nd ACM SIGKDD International Conference on Knowledge Discovery and Data Mining (ACM), pp 785–794.

31. J. M. Colby, B. A. Krantz, Comparative study of nanopore phenylalanine clamp variants reveals unique peptide biosensing and classification properties. (2025).

32. A. F. Kintzer et al., The protective antigen component of anthrax toxin forms functional octameric complexes. J. Mol. Biol. 392, 614–629 (2009).

33. A. Bernard, M. Payton, Fermentation and growth of Escherichia coli for optimal protein production. J. E. Coligan, B. M. Dunn, H. L. Plough, D. W. Speicher, P. T. Wingfield, Eds., Current Protocols in Protein Science (John Wiley & Sons, Inc., New York, 1995), vol. 5.3, pp. 1–18.

34. K. L. Thoren, E. J. Worden, J. M. Yassif, B. A. Krantz, Lethal factor unfolding is the most force-dependent step of anthrax toxin translocation. Proc. Natl Acad. Sci. U.S.A. 106, 21555–21560 (2009).

35. M. Abadi, et al. (2015) TensorFlow: Large-Scale Machine Learning on Heterogeneous Distributed Systems.

36. P. Remy, Temporal convolutional networks for keras. GitHub repository (2020).

37. E. F. Pettersen et al., UCSF Chimera—a visualization system for exploratory research and analysis. J. Comput. Chem. 25, 1605–1612 (2004).

